# Tissue-specificity of gene expression in the *Ciona* embryo is subtly bimodal

**DOI:** 10.64898/2025.12.30.697136

**Authors:** Michael T. Veeman, Kendal R. Palmgren, Cathryn L. Haas

## Abstract

Expressed genes potentially fall into two distinct categories: ‘tissue-specific’ genes expressed in a subset of cell types that carry out the distinctive functions of those cells; and ‘housekeeping’ genes that are broadly expressed across all cell types and carry out the basic functions of life. It is unclear, however, whether these are actually two distinct classes or whether they represent an intuitive but false dichotomy imposed upon gene expression patterns that vary widely and continuously in how specific they are to particular tissues. We address this question using a high-coverage, whole-embryo single cell RNAseq atlas of the model invertebrate chordate *Ciona robusta*. There is a major complication in that quantitative measures of tissue-specificity such as the Tau and Gini metrics show a strong negative correlation with expression level. We show here that this correlation is the result of sampling error and not a fundamental biological relationship. Raw Tau scores are bimodal, but this is largely an artifact of Tau’s sigmoidal relationship with expression level for uniformly expressed genes. Simulations and statistical analyses indicate that the Tau metric is badly confounded by expression level for ubiquitously and/or weakly expressed genes but is relatively accurate for genes that have statistical evidence of differential expression. The distribution of tissue-specificity scores for these differentially expressed genes is broad and flat, spanning from near-binary to near-uniform. While only subtly bimodal, ubiquitously expressed and tissue-specific genes are clearly distinguishable, especially at higher expression levels. We explore the use of pseudocounts to shrink the high tissue-specificity scores of weakly expressed genes and find that they are effective at separating tissue-specific from ubiquitously expressed genes but distort rankings of tissue-specificity. Gene ontology code analysis indicates that the most tissue-specific genes are strongly enriched for predicted roles as transcriptional regulators and tissue-specific effector molecules, whereas the most ubiquitously expressed genes are enriched for predicted housekeeping functions. We conclude that the ‘tissue-specific’ vs ‘housekeeping’ dichotomy is meaningful in the *Ciona* embryo despite the broad range of Tau scores for tissue-specific genes. These findings provide a framework for formally assessing the tissue-specificity of gene expression across a broad range of taxa and developmental stages.

## Introduction

Embryonic development is known to involve transcription factors, signaling molecules and effector proteins expressed under fine spatiotemporal control in specific cell types and developmental stages. These interact through complex regulatory networks to establish new cell types with their own characteristic gene expression profiles and temporal dynamics. By contrast, fundamental biological processes such as cellular metabolism and genome maintenance tend to be controlled by the products of genes that are ubiquitously expressed in all cell types. This distinction between so-called ‘tissue-specific’ and ‘housekeeping’ genes is uncontroversial in many ways. There are many genes known to have very specific gene expression patterns and others that are very broadly expressed, and there are well established correlations between these categories and different cellular functions [1–9].

That said, it is surprisingly unclear how the tissue-specificity of gene expression is distributed across the full set of expressed genes in any species. Large scale *in situ* hybridization screens have been effective at identifying candidate developmental regulators with very specific expression patterns, but ‘messier’ expression patterns tend to get overlooked [10–13]. Traditional *in situs* are not quantitative and often focus on a biased set of genes. Transcriptomic approaches such as microarrays and bulk RNAseq provide a more quantitative and genome-wide view of gene expression, but typically lack spatial and cell type-specific resolution. Single cell RNAseq (scRNAseq) provides a more powerful approach to quantify the tissue-specificity of gene expression with both genome-wide and organism-wide scope, but the sparse, noisy statistics of transcript counts from single cells poses major analytical challenges. Most scRNAseq studies have focused on the identification of distinct cell types and the inference of key regulators and network interactions driving cell state transitions [14–23]. These studies typically focus on genes that are quite tissue-specific in their expression patterns.

There has been extensive work on the identification of Differentially Expressed Genes (DEGs) in bulk and single cell RNAseq data, but this has typically focused on identifying the most informative marker genes to cluster cells into distinct cell types [24–26] and on statistical testing for differential expression between cell types or experimental conditions [27–30]. There has also been work on the identification of Stably Expressed Genes (SEGs) or Housekeeping Genes (HKs) [1,2,4,6–9,31], though this has most often focused on the identification of a small set of particularly uniform genes to be used in normalization strategies. Note that the ‘Housekeeping’ terminology can be used in different ways. It often refers simply to uniform expression across cell types or conditions, but a more rigorous set of criteria also includes predicted functions in fundamental biological processes, essentiality in cell or organismal viability screens, and evolutionary conservation across species [8]. Despite this interest in both tissue-specific and housekeeping genes, there has been little attention given to the systems-level question of whether the tissue-specificity of gene expression is truly bimodal, with distinct sets of tissue-specific and ubiquitously expressed genes, or whether there is instead a continuum of tissue-specificity from very high to very low. The tissue-specific versus housekeeping concept is longstanding and intuitive, but it could be an artificial dichotomy that scientists have imposed on patterns of gene expression that are subtler and more complex. It is noteworthy that the identification of marker genes defining distinct cell types in scRNAseq data has proven to be a challenging topic with numerous competing approaches. A recent benchmarking study compared 59 different methods for scRNAseq marker gene identification [26] and best practices remain controversial. This would arguably not be such a difficult challenge if the tissue-specificity of gene expression was indeed strongly bimodal.

Differences in gene expression between two cell types or between other conditions are commonly represented as log-transformed fold changes. Other metrics have been developed to quantify the tissue-specificity of gene expression across more than two conditions or cell types. Kryuchkova-Mostacci and Robinson-Rechavi benchmarked several and found that the Tau [32] and Gini [31] metrics were most robust to different normalization methods and dataset depths, and correlated well across different profiling methods [33]. Tau and Gini both use a 0-1 scale where 0 represents completely uniform expression across cell types and 1 represents expression that is entirely restricted to a single cell type.

The question of whether the tissue-specificity of gene expression is demonstrably bimodal is newly addressable given the growing number of organisms that have been deeply sampled by whole-organism scRNAseq [14–16,19,23,34–38]. This is particularly true for the embryonic stages of model organisms that have a modest number of well characterized cell types. The invertebrate chordate *Ciona robusta* is noteworthy in this context. *Ciona* embryos have chordate anatomy including a notochord and hollow, dorsal neural tube, but in a particularly compact embryo with a small number of well-characterized cell types [39,40]. These develop through invariant cell lineages that have been intensively studied, such there are strong prior expectations on the number of distinct cell types expected during early embryonic stages. There have been several *Ciona* scRNAseq atlas papers [38,41–46]. Hotta stage 12 (mid-gastrula) is of particular interest as it is early in embryonic development (∼5.7 hours post fertilization at 18°C) [47] but nearly all blastomeres are already restricted to a single tissue type and the initial anterior-posterior and mediolateral patterning of the neural plate has already been established. Our group’s dropSEQ analysis of the mid-gastrula stage identified 28 distinct cell types representing the majority of those expected at this timepoint [46]. As with many scRNAseq methods, dropSEQ uses Unique Molecular Identifier (UMI) barcodes to help correct for PCR amplification biases [48]. scRNAseq data is sparse and noisy at the level of individual cells but UMI counts can be aggregated by inferred cell type into ‘pseudobulked’ representations with greatly improved quantitative properties. Our goal here is to use pseudobulked data from the mid-gastrula *Ciona* embryo to address whether the tissue-specificity of embryonic gene expression is bimodal when assessed across the entire embryo at this landmark stage.

## Results

### Tissue-specificity metrics are negatively correlated with expression level

We reclustered and reannotated the mid-gastrula dataset from Winkley et al [46] in Scanpy [25] to provide a more reproducible analysis using more recent methodology (Fig 1A). There are some finer subdivisions of the neural plate that we failed to detect in our previous analysis, and we speculated that these might be resolvable with newer analysis tools. Despite extensive exploration of different normalization strategies, marker gene selection methods and clustering approaches, we did not detect any additional cell types with high confidence. We found our previous distinction between sibling blastomeres A9.30 and A9.29 to be marginal and kept them as a single A-line column III cell type for a total of 27 distinct cell types.

**Figure 1.**
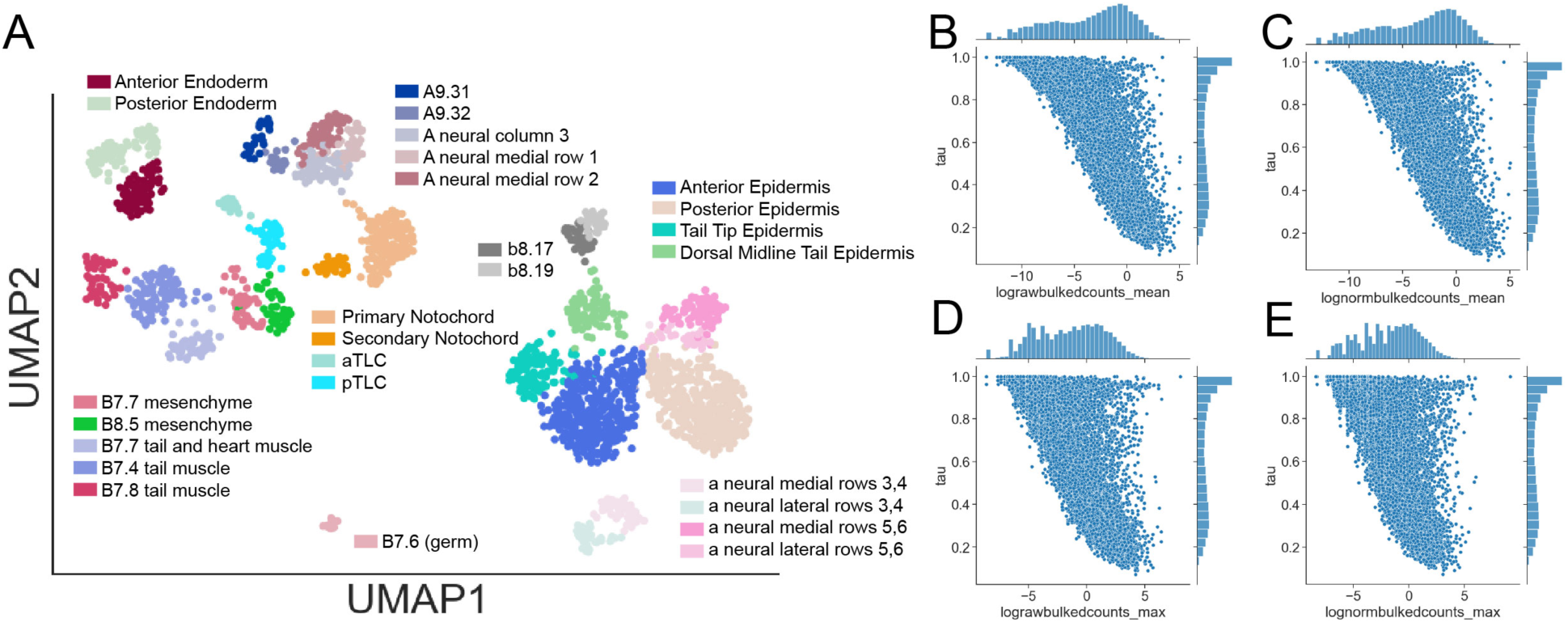
Tau is negatively correlated with read counts. 1A) UMAP plot for the mid-gastrula *Ciona* embryo. Each data point represents a single transcriptionally profiled cell. Points that are close together have similar gene expression profiles. Labelled colors mark the 27 inferred cell types. 1B-E) Scatterplot showing the relationship between expression level and the Tau metric of tissue specificity using 4 different read count metrics. 1B and 1C show mean pseudobulked expression across all 27 cell types. 1D and 1E show the pseudobulked expression in the cell type with highest expression for each gene. 1B and D show raw pseudobulked counts (mean per cell) whereas 1C and 1E are normalized to median total counts per cell type. All read count metrics are log2 transformed for visualization purposes.

The dataset includes 2185 profiled cells passing quality control tests. Cells profiled per cell type range from a low of 14 in the B7.6 germ cell lineage to a high of 397 in posterior epidermis (mean 80.9, median 48). We bulked the UMI counts from each cell type and treated them as either raw bulked counts or performed a simple normalization to median sequencing depth. We further reduced these to a single number per gene model by taking either the mean or the maximum across all 27 cell types.

For each gene with at least at one UMI detected, we calculated the Tau metric based on UMI counts per cell type normalized to median total UMI count per cell type. Tau shows a strong negative correlation with UMI counts that is evident using both raw and normalized counts and using both the mean across cell types as well as the maximum across cell types (Fig. 1B-E). A density plot shows that most transcripts fall roughly along the diagonal (Fig. 2A’ vs Fig. 2A), with the remaining data points falling above that line of density.

**Figure 2.**
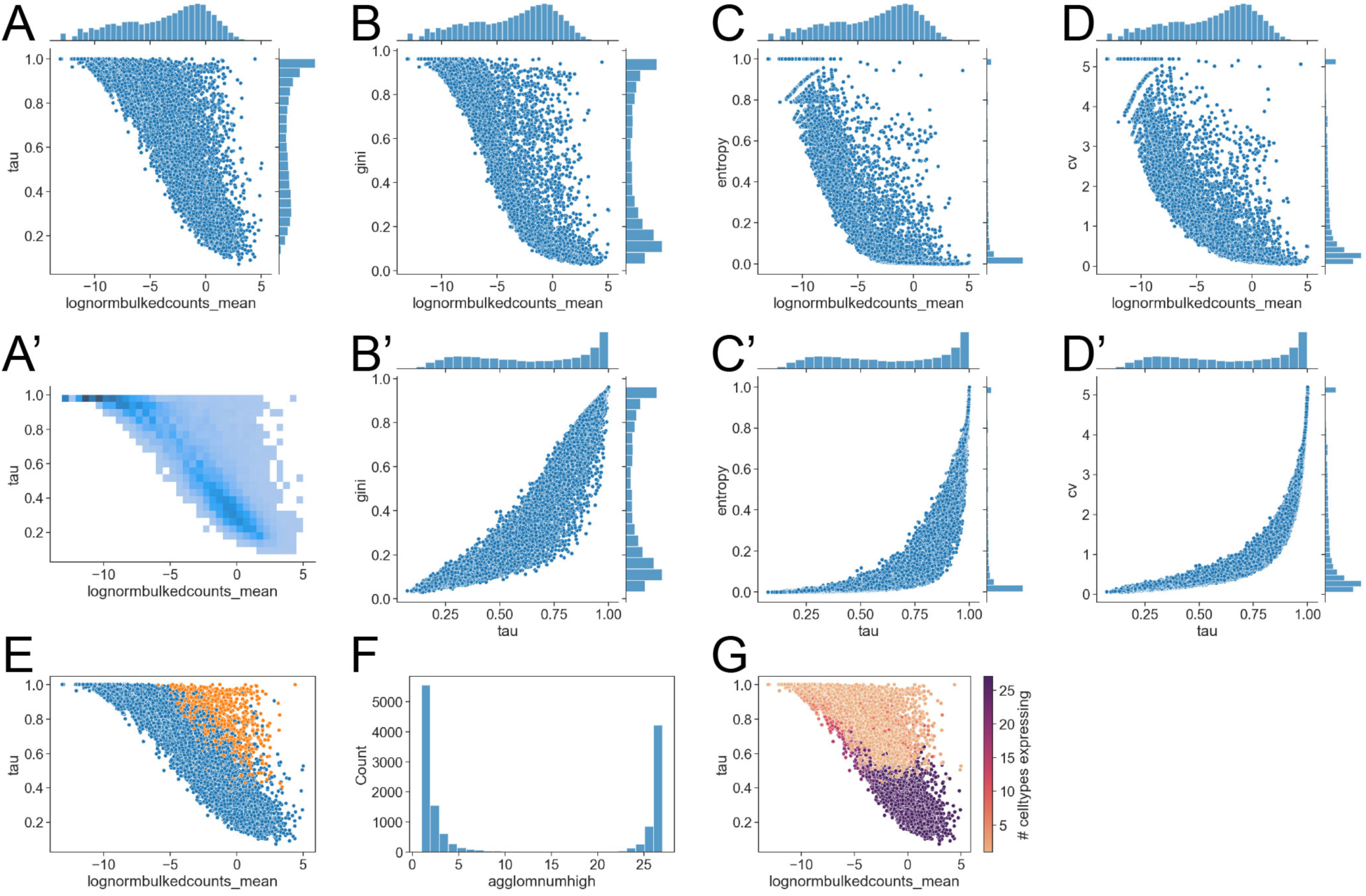
Comparing tissue-specificity metrics. A) Scatter plot of the relationship between log2 transformed normalized mean counts and the Tau tissue-specific expression metric. A’) Density plot (2D histogram) of the scatterplot in A). B) Scatter plot of the relationship between log2 transformed normalized mean counts and the Gini tissue-specific expression metric. C) Scatter plot of the relationship between log2 transformed normalized mean counts and the normalized entropy index. D) Scatter plot of the relationship between log2 transformed normalized mean counts and the Coefficient of Variation (CV). B’,C’D’) Scatter plots showing the correlation between Tau and Gini, normalized entropy and CV respectively.E) Tau/read count plot with Scanpy tissue-specific marker genes (details in methods) highlighted in red. F) Histogram distribution of the number of cell types expressing each gene as inferred by anchored agglomerative clustering (details in Methods). G) The number of expressing cell types visualized as a pseudocolor on the read count/Tau curve.

We also calculated three other metrics of tissue specificity: the Gini coefficient, a 0-1 index based on normalized entropy, and the coefficient of variation (CV) (details in Methods). These also showed pronounced negative correlations with UMI counts (Fig. 2B-D). The curves vary, but the relationship seems at least somewhat sigmoidal for the 4 tissue-specificity metrics tested. All 4 metrics are positively correlated with one another (Fig. 2B’-D’). The relationships between these are broadly monotonic but not linear. As compared to Tau, Gini and the other two metrics group stretch out the differences between the most tissue-specific genes and clump the less tissue-specific genes closer together. We chose to focus on Tau for subsequent analyses as it thus seemed well suited to resolving differences between genes of moderate to low tissue-specificity.

As an initial exploration of the 2D space defined by expression level (normalized UMI counts) and Tau, we labelled the data points based on whether they were called as marker genes based on ‘one versus all’ pseudobulked Wilcoxon rank sum test comparisons in Scanpy (Fig. 2E). The genes with evidence of differential expression across cell types are all in the upper right area of the plot (high tau, high expression), above the sigmoidal curve of greatest density. We also used a threshold-free anchored agglomerative clustering approach (details in Methods) to estimate the number of cell types that strongly express each gene. This metric was sharply bimodal, with most expressed genes strongly expressed in either <=5 cell types or >=23 cell types (Fig. 2F). The putative ubiquitously expressed genes are all at the bottom right of the expression/Tau plot with Tau less than ∼0.5 (Fig. 2G).

The distribution of Tau and other tissue-specificity metrics is distinctly bimodal (right marginal histograms in Fig. 1B-D, Fig. 2A-D). It is important to note, however, that most genes with high tissue specificity scores are weakly expressed, often with only a few counts in one or two cell types. For weakly expressed genes, it is difficult to know whether they are indeed specific to those cell types, or whether they might be more ubiquitously expressed but only marginally detectable and thus only observed in a few cell types. It is similarly unclear whether the bimodal distribution of raw tau scores represents two true modes of tissue-specificity or whether it might be an artifact of an underlying sigmoidal relationship between expression level and tissue-specificity. A statistical framework is therefore needed for assessing the effects of sampling error and detection limits on the tau/expression level relationship.

### Sampling error creates a sigmoidal relationship between tau and expression level for uniformly expressed genes

Differential expression testing between pairs of cell types has been intensively studied. There is a well-known negative correlation between expression level and log fold change between paired samples that reflects statistical sampling error [28]. We hypothesized that the same would be true for the Tau metric. To test this, we performed Monte Carlo simulations to predict the Tau values that would be observed across a broad range of expression levels for genes expressed uniformly across all tissues. We repeated these simulations using different underlying statistical assumptions, different numbers of cell types and different numbers of cells captured per cell type (Fig. 3).

**Figure 3.**
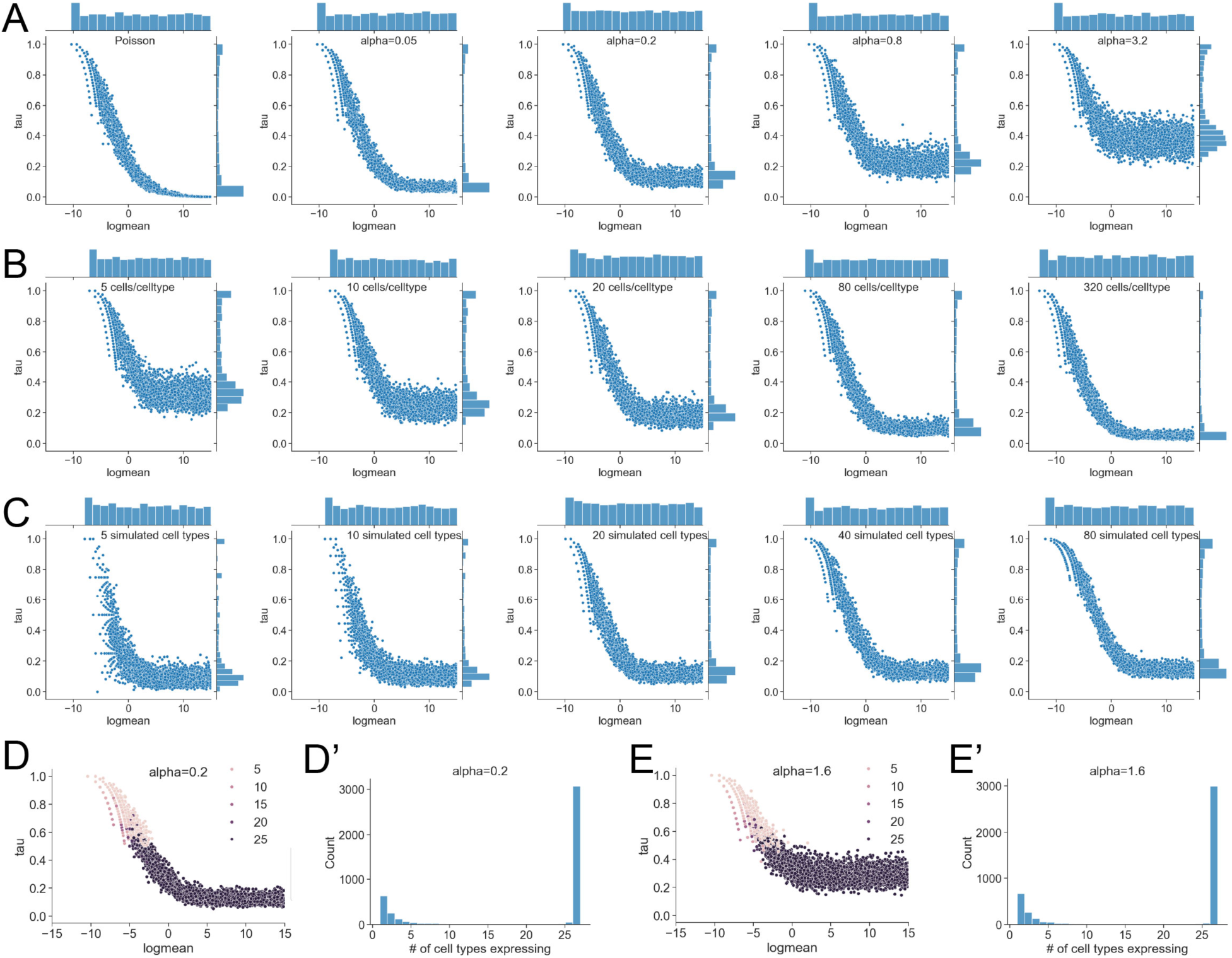
Monte Carlo simulation of uniform expression. A,B,C) Tau/read count curves for simulated data under the assumption of uniform expression across cell types. Each plot is based on 5,000 simulated genes drawn randomly from a broad uniform distribution of different expression levels. Each tau value is based on simulated cell-level read counts in a number of simulated cells and simulated cell types by drawing from either the Poisson or Negative Binomial distribution using the same expression level parameter for each cell type. Unless otherwise indicated each set of simulations uses a negative binomial model with 27 cell types, 50 cells per cell type and alpha=0.2. The plots in A) start with a Poisson model at the left and then progress through a series of negative binomial models with increasing alpha overdispersion factors. The plots in B) vary the number of simulated cells per cell type while holding other factors constant. The plots in C) vary the number of simulated cell types while holding other factors constant. The labeled scatterplots in D and E and the histograms in D’ and E’ show the number of cell types inferred to be expressing each gene based on the anchored agglomerative clustering criterion.

We first simulated the Tau/read count relationship using a Poisson model. This is a standard model for simple simulations of count-based data, though it makes the non-trivial assumption that mean and variance are equal across all expression levels [49]. The Poisson distribution has a single parameter, lambda, that describes both the mean number of counts expected per cell and the variance of those counts. For each simulated gene, we randomly selected a lambda value from a broad uniform distribution and then applied it to all the simulated cells in all the simulated cell types. We then bulked these results by cell type and calculated Tau. Repeated across thousands of simulated genes from a broad range of expression levels, this produces a distinctly sigmoidal curve in which Tau decreases from ∼1 at the threshold of detectability towards ∼0 at high read counts (Fig. 3A, leftmost panel). Notably, the sigmoidal shape of this relationship gives a bimodal distribution of tau despite the uniform lambda parameter across cell types.

Actual RNAseq data tends to be overdispersed with variance greater than the mean. This may commonly reflect latent cell types or states that are grouped together in bulked, pseudobulked, or single cell analyses, but it may also reflect other phenomena such as bursty transcription. Negative binomial models are widely used to represent overdispersed count-based data [50]. The negative binomial distribution can be viewed as a mixture of Poisson models with variable lambda parameters taken from a Gamma distribution. For expression analysis it is usually parametrized as mu (rate parameter similar to Poisson lambda) and alpha (overdispersion parameter). We repeated the Monte Carlo simulations using a negative binomial model and a range of different alpha values (Fig. 3A). As with the Poisson simulations, we kept the expected expression level the same across all cells and cell types for each simulated gene by using the same mu, but randomly selected from a broad range of mus for each simulated gene. Tau and expression level still show a negative sigmoidal relationship, but instead of asymptotically approaching 0 at high expression levels, the relationship plateaus at a higher Tau level (Fig. 3A). The height of this plateau depends on the overdispersion parameter alpha, with higher alpha giving a higher and noisier plateau.

We further explored how the number of cells captured per cell type (Fig. 3B) and the number of distinct cell types (Fig. 3C) influence the relationship between tissue specificity and expression level for uniformly expressed genes. These again had complex effects on the Tau/read count relationship though it remained sigmoidal in all cases. An increasing number of cells assessed per cell type makes the right plateau lower and less noisy (Fig. 3B). An increasing number of different cell types makes the left shoulder of the sigmoidal curve more distinct and has subtler effects on the right plateau (Fig. 3C). We also calculated the number of expressing cell types using the agglomerative clustering approach from Fig. 2F,G and found it to be strongly bimodal with a rapid transition at increasing expression levels between expression in just one or a few simulated cell types to expression in all or nearly all cell types (Fig. 3D,D’,E,E’).

### Monte Carlo simulation of Tau for differentially expressed genes

These simulations indicate that statistical sampling error is sufficient to explain the decreasing sigmoidal relationship between tissue-specificity and expression level for genes that are uniformly expressed across cell types. It is unclear, however, how sampling error affects Tau for genes that are differentially expressed across cell types. We thus performed additional Monte Carlo simulations to explore the scenario where there is a base level of expression across all cell types but one cell type expresses at a higher level. The fold change in mu between the single high cell type and the remaining low cell types controls the theoretical Tau that would be expected assuming perfect sampling. We again simulated how observed Tau would vary across a broad range of read counts for different theoretical Taus (Fig. 4). These simulations were implemented using a negative binomial model with 3 different alpha values: near-Poisson (Fig. 4A); 0.2 (Fig. 4B); and 0.5 (Fig. 4C).

**Figure 4.**
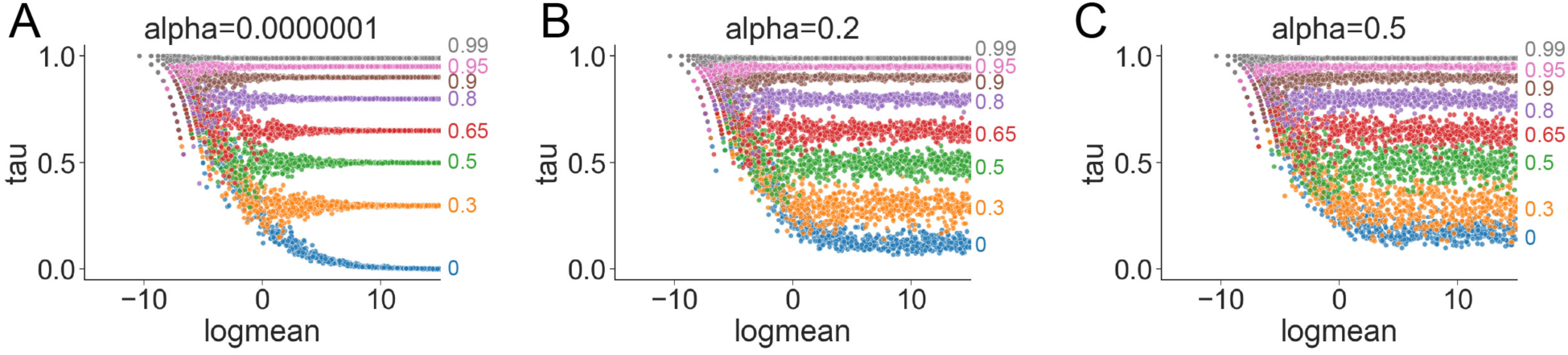
Monte Carlo simulation of non-uniform expression. A,B,C) Tau/read count curves for simulated data assuming differential expression between one cell type and the remaining cell types. The log fold changes between high and low cell types were chosen to give the indicated theoretical Tau levels assuming perfect sampling. The three models differ in the overdispersion parameter alpha which varies from A) near-Poissonian to B) low, and C) moderate.

The resulting curves are similar to those for uniformly expressed genes at lower read counts but diverge at higher read counts to plateau close to the theoretical Tau level. This divergence is rapid with the observed Tau diverging from the uniform expression curve to plateau over a relatively small change in expression level. It then remains flat at close to the expected Tau level at higher expression levels. While these simulations only test one type of differential expression profile, they suggest that observed Taus are likely to be relatively accurate for genes with enough read counts to detect differential expression. The same general relationships are seen for all alpha values tested, but higher alpha values increase noise in the plateau (decreased precision) for all theoretical Tau values and increase the height of the baseline plateau for uniform expression (Fig. 4).

### Hypothesis testing framework for Tau being higher than predicted based on uniform expression

We next developed a Monte Carlo approach to identify which gene-specific Tau values from the mid-gastrula *Ciona* dataset are higher than predicted under the assumption of uniform expression across all cell types. We used 4 different read count models for these analyses: a Poisson model; a negative binomial model with a common alpha value derived from the mean/variance relationship across all genes and cell types, and two negative binomial models with gene-specific alpha values fitted using different underlying assumptions. Parameter values for each gene were estimated by fitting a Generalized Linear Model (GLM) to the raw count data for each captured cell using sequencing depth as an offset. These parameters were then used in Monte Carlo simulations based on the actual number of cells captured for each cell type in the mid-gastrula *Ciona* dataset and the actual sequencing depth for each cell, but under the assumption of a uniform lambda or mu parameter for each cell type. For each model we performed 25,000 Monte Carlo replicates for each gene and used these to calculate a median predicted Tau for uniform expression as well as a p-value for the observed Tau being higher than predicted based on uniform expression across cell types. These p-values were then adjusted using the Benjamini-Hochberg false discovery rate correction for multiple comparisons [51].

Fig. 5A shows the mean/variance relationship for all expressed genes calculated across all cells without regard to inferred cell type. Fig. 5B shows the mean/variance relationship for all expressed genes calculated separately for each cell type. Both show evidence of overdispersion with variances being consistently higher than the mean at higher expression levels. For the standard negative binomial model, variance relates to the mean according to v=µ+⍺µ2. For the common alpha model, we calculated a global estimate of a single alpha for all genes by fitting this polynomial to the mean/variance data and solving for alpha by nonlinear least squares. Calculated without regard to inferred cell type this gives a common alpha of 0.65. Using means and variances calculated separately for each cell type, this gives a common alpha of 0.38. We used the latter value for the common alpha Monte Carlo model as we are simulating uniform expression across cell types and differential expression across cell types is likely the main reason for the difference between these two estimates.

**Figure 5.**
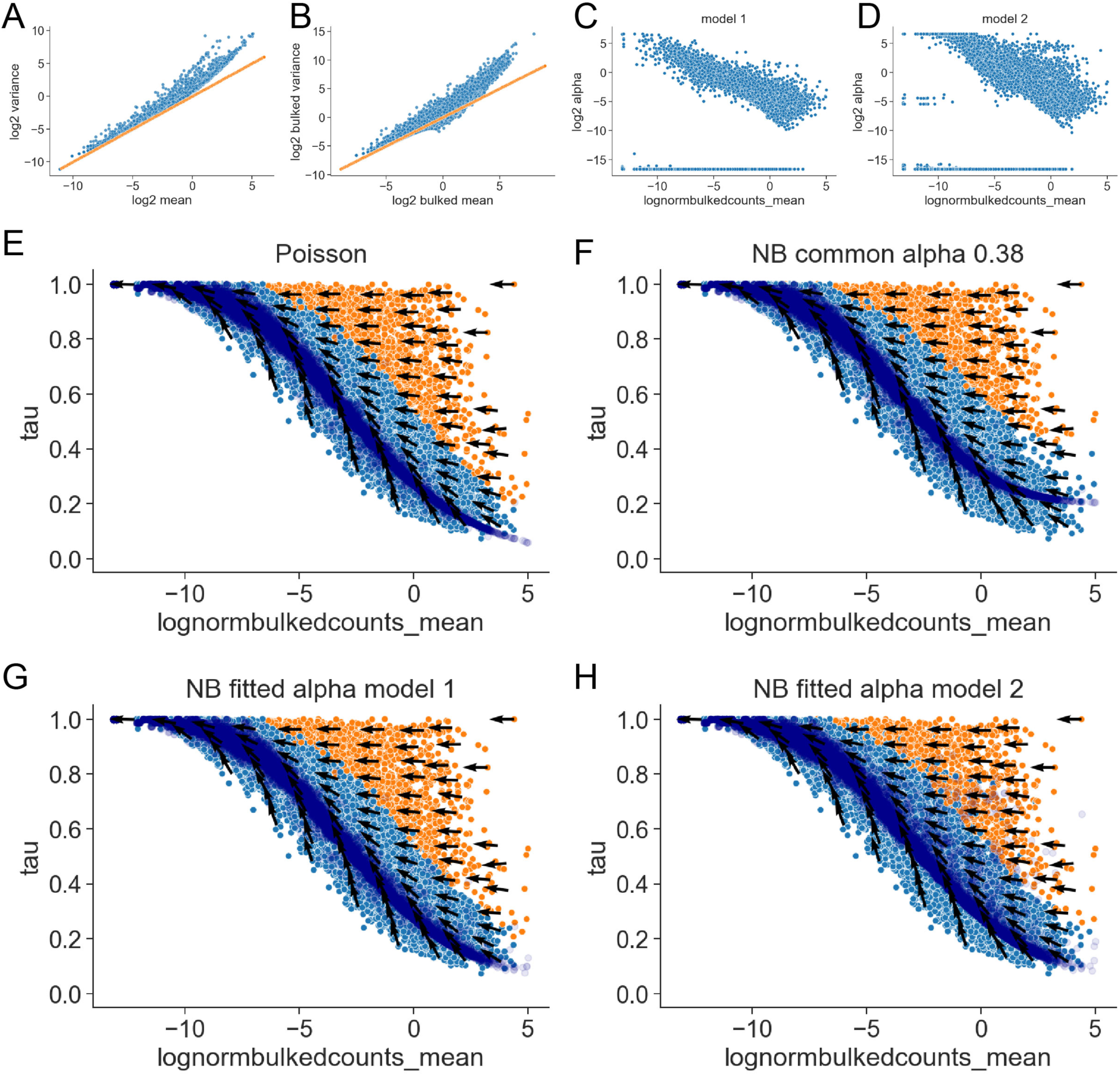
Hypothesis testing for whether Tau is greater than predicted for uniform expression. A) Mean/variance plot for all cells (not aggregated by cell type). B) Mean/variance plot (aggregated by cell type). Orange lines in A and B indicate the y=x diagonal. C) Scatter plot of the alpha overdispersion parameter for each gene as a function of read counts. Alpha estimated under model 1 assumptions that allow a different expression level parameter for each cell type. D) Scatter plot of the alpha overdispersion parameter for each gene as a function of read counts. Alpha estimated under model 2 assumptions with a common expression level parameter for all cell types. E-H) Tau/read count plots showing p-values for Tau being greater than predicted under the assumption of uniform expression across cell types. Genes with Tau p-values <0.05 after correction for multiple comparison are shown in orange. Genes with non-significant Tau p-values are in pale blue. The dark blue overlay indicates median Tau values predicted for each gene if they were uniformly expressed across all cell types. The streamline arrows indicate locally averaged predicted changes in Tau for a 50% decrease in expression. E-H) differ only in the model assumptions used to calculate p-values: E) Poisson; F) Negative Binomial, alpha=0.38 for all genes; G) Negative Binomial with alpha fitted for each gene under model 1 assumptions; H) Negative Binomial with alpha fitted for each gene under model 2 assumptions.

For gene-specific alpha model 1, we fitted an initial GLM using inferred cell type as a categorical variable to estimate an alpha value for each gene and then used that alpha to estimate mu under the assumption of uniform expression. For gene-specific alpha model 2, we fitted alpha and mu naively without regard to inferred cell type. The latter approach is likely to overestimate alpha for differentially expressed genes as it fits a single mu across all cell types. This can be viewed, however, as an upper estimate on the likely range of alpha for uniform expression. Fig. 5C and D show the relationship between expression level and alpha for both gene-specific alpha models. In both cases, there is a negative correlation with more strongly expressed genes having lower alpha values. As predicted, the alpha values fitted for model 2 are higher for many genes, but in both cases, many genes have very low alpha values consistent with near-Poisson expectations. The negative correlation between expression level and overdispersion is not well understood but has been seen in other studies and is not an unusual feature of this dataset [28,50].

Fig. 5E-H show observed Tau vs read count scatterplots for the 4 different models. The dark blue overlay shows the simulated Tau predicted for each gene if it was uniformly expressed across cell types. Gene models with statistical support for observed Tau being higher than predicted under the assumption of uniform expression are highlighted in orange (multiple comparison corrected p-value <0.05). The arrows represent predicted changes in Tau based on a 50% decrease in expression level. These are averaged locally for display purposes. For all 4 models, the dark blue curves of simulated Tau values for uniform expression are reasonably well matched to the distribution of observed Tau values shown in pale blue for genes with non-significant Tau p-values. The streamline vectors indicating the predicted change in Tau for a 50% decrease in expression are relatively flat for the genes with significant Tau p-values but angle sharply towards the simulated uniform Tau medians for the genes with non-significant p-values. This again suggests that the observed Tau values are meaningful for genes that are differentially expressed across cell types but badly confounded by expression level for genes that are uniformly expressed in all cell types.

While they are not identical, the 4 models identify a relatively congruent set of genes as having Tau values higher than predicted based on sampling error alone. This is partly because the overexpression parameter is relatively unimportant for the many genes expressed at less than 1 mean UMI count per cell. For the more strongly expressed genes, the modest level of alpha=0.38 under the common alpha model and the negative correlation between alpha and read counts for the two fitted alpha models are such that the negative binomial models are not grossly different from the Poisson model or from each other. We favor the negative binomial fitted alpha model 1, as it has strong theoretical underpinnings and the distribution of simulated uniform expression Tau values seems closest to the observed data for the non-differentially but strongly expressed genes.

### Pseudocounts have complex effects on the expression level/Tau relationship

In parallel to developing this statistical framework for understanding how read depth influences observed Tau values, we also explored pseudocount addition as a simple strategy for shrinking Tau values for weakly expressed genes. While not appropriate for approaches based on explicitly modeling raw read counts, count-based data is commonly transformed by adding a constant pseudocount followed by log transformation, which substantially normalizes the transformed expression levels. Differential expression testing methods based on simple log pseudocount transformations have been shown to perform surprisingly well compared to more elaborate methods [26]. The pseudocount mitigates the problem of division by zero for genes with no reads and this transformation also helps control the variance across genes with different expression levels. It additionally helps to shrink log fold changes for weakly expressed genes, with the extent of this shrinkage depending on the magnitude of the pseudocount.

Figure 6A shows a series of Tau/read count plots where Tau was calculated after the addition of progressively higher pseudocounts. As shown, pseudocounts bend the left side of the curve downwards, with the extent of the bending depending on the size of the pseudocount. With high pseudocounts, the relationship between expression level and Tau becomes positively correlated as the only genes that have not had tau shrunk to near 0 are the most strongly and differentially expressed.

**Figure 6.**
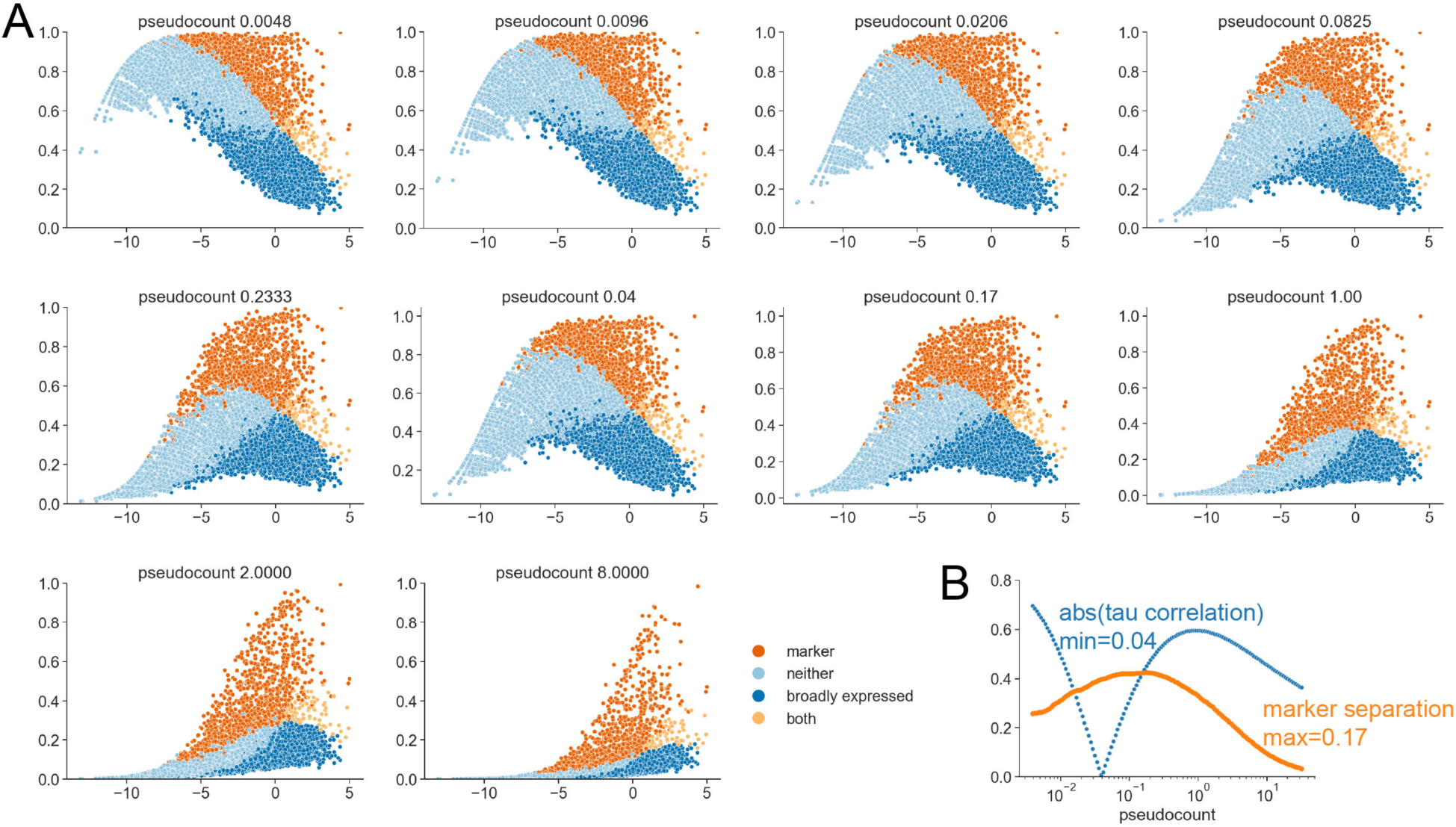
Pseudocount addition distorts the Tau/read count relationship. A) Scatterplots of Tau as a function of log2 normalized mean counts (mean across all cell types). Tau (y-axis) is calculated after the addition of the indicated pseudocount. Read counts (x-axis) are the original value prior to pseudocount addition. ‘marker’ labels are based on multiple comparison corrected Tau p-value<0.05 under the negative binomial fitted alpha model 1 as in Fig. 5G. ‘broadly expressed’ labels are based on the anchored agglomerative high/low clustering criterion as in Fig. 2G. B) Cost function plots for pseudocount optimization. The blue line shows the absolute value of the Pearson correlation between Tau and read counts as a function of the pseudocount. The orange line shows the difference between the median Tau of markers genes and the median Tau of all other expressed genes as a function of the pseudocount.

We reasoned that explicit consideration of the relationship between expression level and observed tissue-specificity could form the basis for a principled choice of pseudocount. One such strategy is to find the pseudocount that minimizes the correlation between expression level and Tau. For this dataset, this occurs with a modest pseudocount of only ∼0.04 mean counts per cell (Fig. 6A,B). Note that with this pseudocount, the expression level/Tau relationship for low tissue-specificity genes is no longer monotonic and now shaped more like an inverted U.

Another strategy is to find the pseudocount that best separates genes inferred to be differentially expressed from those that are not. Using the genes with significant Tau p-values under negative binomial fitted alpha model 1 as markers, the pseudocount that gives the biggest difference between the marker gene median Tau and the non-marker gene median Tau is ∼0.17 (Fig. 6A,B). Note that this high a pseudocount bends the curve down even for high Tau genes where the raw Tau score is likely to be a relatively accurate estimate. It does a good job of separating transcripts with strong evidence of differential expression from those that lack such evidence, but it badly distorts rankings of tissue-specificity within the more tissue-specific genes.

The arbitrary but commonly used pseudocount of 1 has even more drastic effects on the expression level versus Tau relationship. The same is likely to be true for most scRNAseq libraries given typical UMI counts, and for bulk RNAseq data sets after typical CP10K or FPKM normalizations. Large pseudocounts are effective for finding strongly expressed tissue-specific marker genes but are not ideal for transcriptome-wide studies of the concept of tissue-specificity itself.

### Surveying the Tau/read count relationship

To tie the abstract Tau metric back to cell-level representations of differential expression, in Fig. 7 we show UMAP plots pseudocolored by expression level for select genes. These were selected algorithmically along two transects of the Tau/read count plot (Fig. 7A). One transect examines genes with very high Tau greater than 0.98 across the observed range of expression levels (Fig. 7B-L, cyan data points in Fig. 7A). The other transect examines genes with high but not extreme expression levels (between 1.5 and 2.5 mean log2 normalized counts per cell in the highest expression cell type) across the full range of Tau values (Fig. 7M-X, purple data points in Fig. 7A).

**Figure 7.**
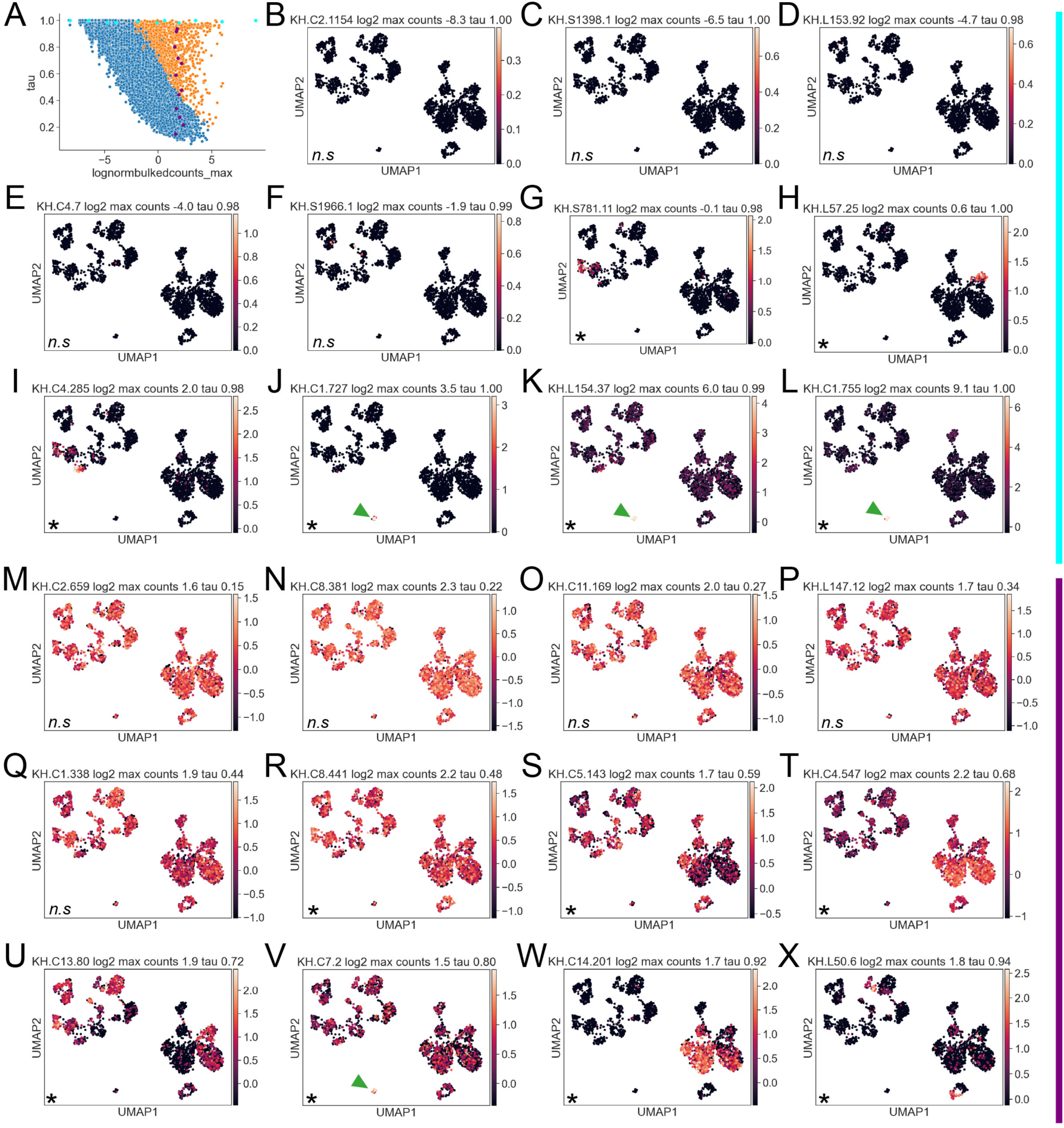
Selected gene-specific UMAP plots along two transects of Tau/read count space. A) Scatterplot of Tau versus log2 normalized mean counts (maximum across all cell types. Orange versus blue labels indicate significant vs non-significant Tau p-values (multiple comparison adjusted p-value<0.05, negative binomial fitted alpha model 1). Cyan and purple labels indicate the selected genes along two orthogonal transects. B-L) The cyan transect explores genes with high Tau >0.98 across the full range of expression levels. M-X) The purple transect explores genes within a zone of moderately high expression across the full range of Tau values. Each UMAP plot is colored to show the expression of the selected gene across all cells and cell types. Note that the color scales vary based on the maximal expression of each gene. Strong expression in the B7.6 germ cell lineage is highlighted with green arrowheads. Each UMAP has an asterisk in the bottom left corner if the Tau p-value is significant or ‘n.s.’ if it is not.

The high Tau transect correlates well visually with the Tau-based differential expression testing. The genes with non-significant Tau p-values are expressed at such low levels that it is difficult to make out where they are expressed. By contrast, the genes with significant Tau p-values all show distinct expression in either one cell type or a small number of cell types. Gene model KH.C1.755 is a noteworthy outlier in this dataset (Fig. 7L). It has a very high Tau of 0.999 but is also the mostly strongly expressed gene in the dataset on the basis of normalized counts per cell in its most strongly expressing cell type. It encodes a maternal factor variously known as Ci-pem or pem-like that becomes segregated to the B7.6 germ cell lineage in a series of asymmetric divisions in the posterior vegetal quadrant of the early *Ciona* embryo. It is known to be functionally important for controlling ascidian posterior vegetal cell fates together with other maternal determinants like the transcription factor KH.C1.727 (Macho-1/Zic-r.a, Fig. 7J) [52–56]. Several of the high expression/high Tau genes have a similar expression pattern (green arrowheads on the UMAP plots).

The orthogonal transect at moderately high expression levels also correlates well with the Tau-based differential expression testing. Differences between cell types are not visually apparent for the low Tau genes with non-significant Tau p-values but become increasingly distinct at higher Tau levels. The genes with Tau values greater than ∼0.9 are expressed relatively cleanly in just one or a few cell types but multiple patterns are seen at intermediate Tau levels. KH.C4.547 (Tau 0.68, Fig. 7T) for example is expressed at some level in all cell types but is considerably stronger in animal hemisphere cell types compared to vegetal cell types. KH.7.2 (Tau 0.8, Fig. 7V) by contrast is strongly expressed in the B7.6 lineage, moderately expressed in the remaining vegetal cell types and more weakly expressed in the animal cell types.

### Evidence for bimodality

Assessing whether the tissue-specificity of gene expression is in any way bimodal is complicated by the sampling error-driven relationship between tissue-specificity metrics like Tau and read counts. This relationship can be partially corrected using pseudocounts (Fig. 6) or by regressing out the component of Tau driven by sampling error under the assumption of uniform expression (not shown). These methods are not compelling here, however, because they also reduce Tau for weakly expressed but convincingly tissue-specific genes where the initial Tau estimate is likely reasonable. Tau is badly confounded by expression level and sequencing depth for genes that are either uniformly expressed or expressed at read counts that are too low to detect subtle differential expression, but it does not appear to need substantial correction for genes where Tau is significantly higher than predicted under the assumption of uniform expression.

A reasonable strategy for assessing the potential bimodality of Tau is thus to assess it within specific ranges of expression as informed by model-driven insights about Tau p-values. Fig. 8A shows the Tau/read count scatterplot using log normalized counts in the maximum expression cell type for each gene. This is preferable compared to the mean across cell types as maximum expression levels are similar for the most strongly expressed tissue-specific genes and the most strongly expressed ubiquitously genes. Fig. 8B-F show histogram distributions of Tau for specific ranges of read count. For the most strongly expressed genes with log2 normalized max counts >2.5, the distribution of Tau is unequivocally bimodal with a distinct low tissue-specificity peak at Tau ∼0.2 and a high expression peak ending at Tau ∼1 (Fig. 8B). For a larger expression range of log2 normalized max counts >0, there is still a normally distributed low peak centered on Tau ∼0.25, but the higher Tau genes are now uniformly distributed across the remaining range of Tau (Fig. 8C). Histogram distributions of Tau across lower read count slices show a progressively higher and broader uniform expression peak and a progressively narrower range of uniformly-distributed high Tau values (Fig. 8D-E). We conclude that the distribution of Tau is bimodal, but only sharply bimodal for the most strongly expressed genes. For most expression levels that are high enough to detect differential expression across cell types, the Tau distribution is arguably best viewed as a mixture of a normal distribution of uniformly expressed genes and a broad uniform distribution of differentially expressed genes.

**Figure 8.**
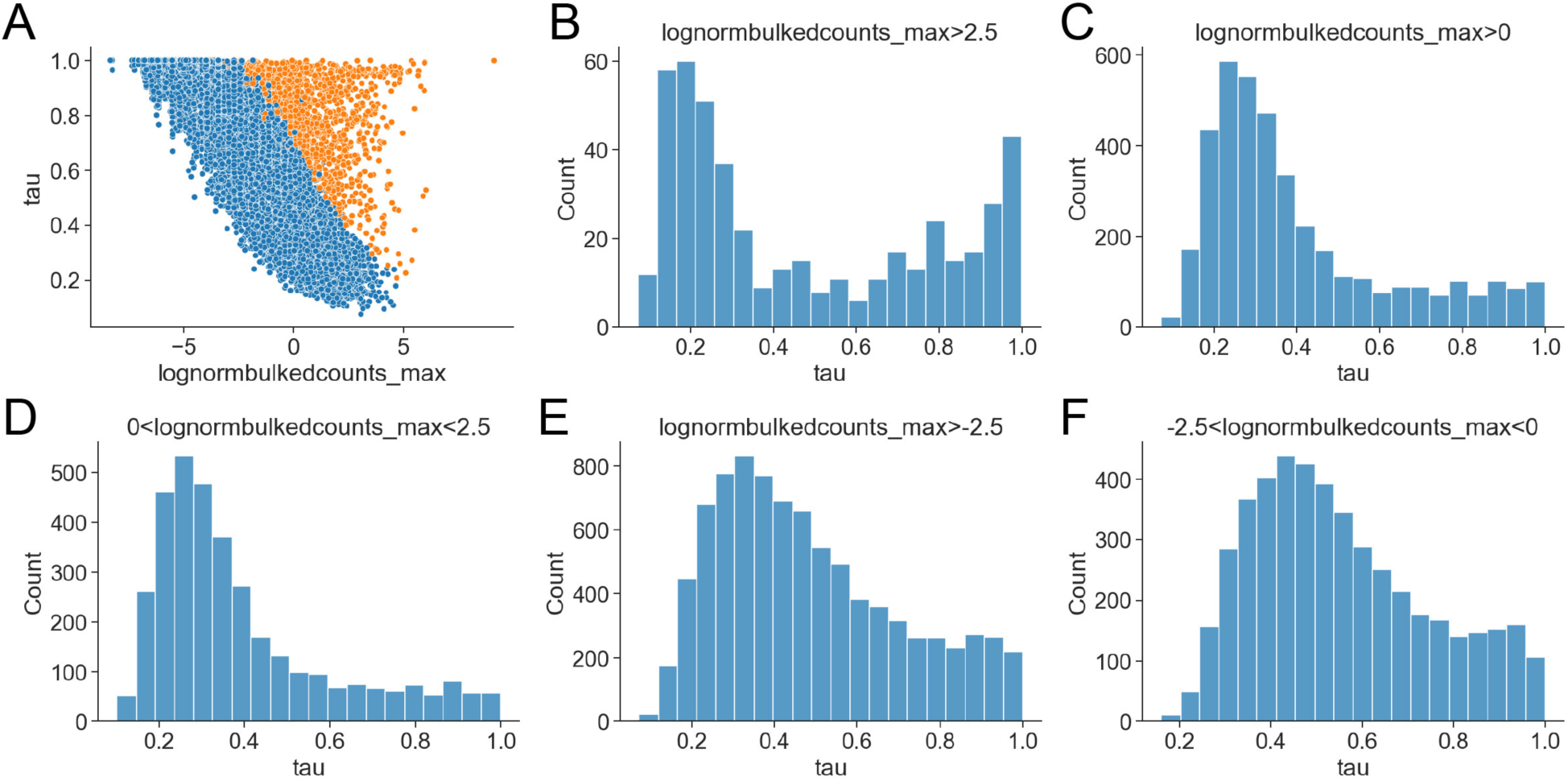
Tau is subtly bimodal. A) Scatterplot of Tau versus log2 normalized mean counts (maximum across all cell types. Orange versus blue labels indicate significant vs non-significant Tau p-values (multiple comparison adjusted p-value<0.05, negative binomial fitted alpha model 1). B-F) Histogram distributions of Tau for the indicated expression level ranges.

### GO Code enrichment supports predicted roles in developmental versus housekeeping functions

To interrogate the potential functions of the distinct gene sets identified, we performed Gene Ontology (GO) code enrichment analysis. We used a preexisting GO annotation of *Ciona* gene models based on homology to vertebrate orthologs [57] and the GOAtools enrichment analysis package [58]. We quantified GO code enrichment for three distinct gene sets in comparison to all genes with detectable expression in this mid-gastrula *Ciona* scRNAseq dataset: all tissue-specific genes with Tau p-value <0.05 after correction for multiple comparisons; the most tissue-specific genes with Tau>0.9 and corrected Tau p-value <0.05, and the set of ubiquitously expressed genes with corrected Tau p-value >0.05 and expression in most or all cell types by the agglomerative clustering criterion. Output tables with full details are in Supplemental Spreadsheets 1-3.

For the set of all genes with a significant Tau p-value, the most enriched GO codes related to transcription factor activity, embryonic development, signaling and morphogenesis. GO codes related to transcriptional regulation were particularly enriched. For the subset of particularly tissue-specific genes with significant Tau values >0.9, similar GO codes were enriched but there were now also many GO codes related to muscle function and contractility. Primary muscle fate is specified early in *Ciona* development and there are many muscle-specific effector genes among the most strongly and differentially expressed transcripts.

The ubiquitously expressed gene set revealed a completely different suite of enriched GO codes. Here the most enriched codes related to basic cellular functions including RNA processing, translation, ribosome biogenesis, vesicle transport and DNA repair.

## Conclusions

### Tissue-specificity is subtly bimodal

Despite Tau being only subtly bimodal, these results suggest that the ‘tissue-specific’ vs ‘housekeeping’ gene dichotomy is meaningful. A proper statistical framework is needed to account for the partial dependence of Tau on expression level, but Tau values higher than predicted based on sampling error under the null hypothesis of uniform expression are readily discernable. Most genes do not have statistical evidence for Tau being greater than predicted based on sampling error alone, and they fall along a roughly sigmoidal curve from Tau ∼1 for the most weakly expressed genes to Tau ∼0.2 for the most strongly expressed. The space above that curve in the upper right diagonal is filled with a broad, flat distribution of tissue-specific genes ranging from near-binary patterns at Tau close to 1 to very subtle non-uniform patterns for genes close to the uniform expression curve. The tissue-specific and ubiquitously expressed gene sets meet expectations in terms of predicted functions as developmental regulators and tissue specific effectors versus housekeeping genes encoding basic cellular functions. The overall distribution of Tau would not be well represented by a mixture of two Gaussians except at the highest expression levels, but it could be represented by a mixture of a Gaussian distribution of uniformly-expressed genes and a uniform distribution of genes that are themselves unevenly expressed across tissues. Ubiquitous and tissue-specific genes are separable and functionally different, but tissue-specific genes vary broadly in how tissue-specific they are. Relatively few are near-binary, and many are quite subtle in how they differ between cell types.

### Caveats

A major caveat to this work is that we have only examined a single stage quite early in the development of one species. Nearly all *Ciona* blastomeres are restricted to a single tissue type at mid-gastrula [59] but they are very early in their trajectories of differentiation. Many endpoint cell types, like specific classes of neuron, have yet to be established and there is little overt, morphological differentiation of morphologically distinct cell types like muscle, notochord, mesenchyme or neurons. Most cell types will still be diverging transcriptionally from one another, but there may be aspects of gene expression that are converging with respect to various common cell types such as muscle, mesenchyme and neural plate that are each derived from multiple different founder cells with distinct early gene expression profiles [38,46]. It is likely that the distribution of tissue-specificity will change over time, and it may become more distinctly bimodal.

Another important caveat is that the *Ciona* mid-gastrula stage is only a few hours after the major onset of zygotic gene expression at the ∼32-cell stage [60]. The timeframe for the decay of maternal transcripts in *Ciona* is not well understood and it is likely that many or all the ubiquitously expressed genes in this dataset are maternally provided and not newly expressed in the embryo. The spectrum of differential expression may also be quite different in organisms like *Ciona* with stereotyped and deeply invariant early lineages as compared to other species with more regulative development, greater developmental stochasticity, and widespread use of morphogen gradients. That said, the fundamentals of how Tau and other tissue-specificity metrics vary as a function of read counts for different degrees of tissue-specificity should be similar across taxa.

### Potential extensions

The quantitative framework developed here could be extended in several ways. The gene-specific estimates of the overdispersion parameter could potentially be improved using DESEQ2-style shrinkage methods [28]. Bayesian or other methods could be used to calculate confidence intervals for Tau based on the general insight that true Tau values for genes with non-significant Tau p-values are unlikely to be much higher than the observed Tau value but could be anywhere lower. New estimators of Tau or related metrics may also be possible that are less biased by expression level and/or incorporate insights based on the distribution of Tau values at higher read counts where the expression level is less problematic. While not explored here, different metrics are potentially needed to detect genes that are broadly expressed but down-regulated in a small number of cell types. There are also interesting questions about the strengths and weaknesses of detecting differential expression in pseudobulked data based on various iterative pairwise comparison strategies (one to all, one to one…) versus approaches like the one used here based on joint assessment across all cell types. On the empirical side, the quantitative framework developed here would benefit from deep replication across multiple biological replicates, scRNAseq methods, timepoints and taxa.

### scRNAseq UMI counts are typically lower than optimal for assessing Tau

The ability to detect differential expression between cell types depends on both expression level and the degree of differential expression. Our simulations suggest that there is an important inflection point at ∼1 mean UMI count/cell where the tau/read count curve starts to flatten out for uniformly expressed genes. The simulations suggest that profiling an increasing number of cells per cell type will be helpful in reducing the height of Tau’s right plateau and decreasing sampling noise, but it may not move this inflection point to the right. Experimental tradeoffs in the number of cells captured and the count depth at which they are profiled should be carefully weighed in experiments designed to assess the full range of tissue-specificity. Current trends to sparsely profile an increasingly large number of cells may be at odds with the importance of high UMI counts per cell in assessing moderate to low tissue-specificity. That said, absolute transcript counts per cell and scRNAseq mRNA capture efficiency impose limits on achievable UMI count depth. The sigmoidal relationship between observed tissue-specificity and count depth should be kept in mind when designing and interpreting scRNAseq studies.

## Methods

### Computational toolkit

All analyses and simulations were performed in Python using the Pandas [61], Numpy [62] and Scipy [63] data science packages. scRNAseq clustering and annotation used the Scanpy and Anndata packages [25]. Statistical tests and simulations were implemented using the Statsmodels package[64]. Most plots were generated using the Seaborn plotting package [65]. Python scripts to replicate this manuscript’s findings and plots are available at github.com/chordmorph/Tissue-Specificity-of-Gene-Expression.

### Scanpy reannotation and reclustering

The processed Seurat object for the mid-gastrula scRNAseq dataset from Winkley et al [46] was downloaded from GEO and converted to .h5ad format using MuDataSeurat [66]. We reclustered in Scanpy starting from raw UMI counts per *Ciona* KH2012 gene model. For clustering, we normalized to CP10k with sc.pp.normalize_total and then log transformed after adding a pseudocount of 1. Mother of origin effects were regressed out using the mother of origin assignments inferred in [46]. Highly variable genes were identified using Pearson Residuals [67]. UMAP projections [68] and Leiden clustering [69] used the top 50 principal components and a local neighborhood of 15. An initial round of Leiden clustering with a resolution parameter of 1.5 identified 20 cell type clusters. Several of these were then subclustered with finer resolution settings based on knowledge of the *Ciona* embryonic lineage and gene expression patterns. Full details are in the analysis script.

An initial set of tissue-specific marker genes was calculated using the Scanpy rank_genes_groups function grouping by final inferred cell types. These calculations used the Wilcoxon method on log(CP10k+1) transformed data but without regressing out the mother of origin effects. Differential expression tests were carried out for each gene in each cell type compared to the union of all other cell types. The marker gene set for Fig. 2E consists of genes with an fdr_bh corrected Wilcoxon p-value of <0.05 for at least one cell type and a log fold change in that comparison of at least 1.

The log(CP10k+1) normalization was only used for this initial clustering and marker gene identification. Subsequent analyses started anew from raw counts.

### Tissue-specificity metrics

Raw counts were pseudobulked by cell type to give mean UMI counts per cell type, total counts per cell type, and the variance in counts per cell type. Most subsequent analyses were based on mean counts per cell type, which were either left raw or lightly normalized by dividing by the total number of counts in that cell type and then multiplying by the median of total counts across all cell types. Functions to calculate Tau, Gini, CV and normalized entropy were implemented in Python. Unless otherwise noted, these metrics were always calculated on the basis of normalized pseudobulked mean counts per cell.

Tau [32] was calculated as 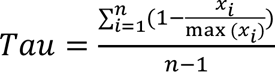 where *xi* is the expression level in cell type *i* and *n* is the total number of cell types.

Gini [31,70] was calculated by numerical integration based on the area between the Lorenz curve (ordered cumulative sum) for each transcript’s gene expression profile and the Lorenz curve for uniform expression.

The base 2 entropy of each gene’s expression profile was calculated with scipy.stats.entropy using the fractional expression in each cell type. For 27 cell types this would have a maximum of log2(27) if all expression is in one cell type. To bring this onto a 0-1 scale with the same directionality as Tau and Gini, we divided each value by log2(27) and subtracted this from 1.

### Anchored agglomerative clustering

We investigated various methods for partitioning expression profiles into On vs Off or High vs Low cell types. Expression profiles with strong expression in some cell types and little to no expression in others can easily be split into high and low groups by rank ordering the expression levels and finding the largest spacing between consecutive expression data points. For one-dimensional transcript profiles across cell types this is equivalent to agglomerative clustering using single linkage. For genes that were broadly expressed, however, this typically led to an arbitrary split between cell types with slightly higher and slightly lower expression. If we added a dummy cell type with no expression, all the real cell types would now be called as On/High. This zero-anchored agglomerative clustering approach produces intuitive splits between On/High and Off/Low across many expression profiles examined. It does not depend on any arbitrary threshold and is not affected by pseudocounts. While slightly different, this method was inspired by the binarization strategy used in [41].

### Monte Carlo simulation (uniform expression)

These simulations were based on simulating read counts on a per cell basis in a specific number of cells across a specific number of tissues. Each read count was simulated by drawing from either the Poisson distribution or the negative binomial distribution. The rate parameter lambda (Poisson) or mu (negative binomial) for each simulated gene was selected by drawing from the uniform distribution with end points chosen to span from below the threshold of detectability to well above the highest mean UMI counts observed in the *Ciona* mid-gastrula scRNAseq dataset. The same rate parameter lambda or mu was used across all the simulated cell types for each simulated gene. After drawing the simulated read counts for each simulated gene, the counts were pseudobulked by cell type. Tau was then calculated based on these simulated pseudobulked mean counts. Default values for the number of simulated genes, number of cells per cell type, number of simulated cell types, and negative binomial overdispersion parameter alpha are given in the Fig. 3 legend.

### Monte Carlo simulation (differential expression)

These simulations used the same general framework as above using the negative binomial model. Instead of having a constant rate parameter per cell we now vary the rate parameter between cell types. We simulated a simple mode of differential expression where it is uniform across most cell types but higher in one. An increased fold change between high and low cell types gives an increased theoretical Tau value. Fold changes were selected to give theoretical Tau values of 0, 0.3, 0.5, 0.65, 0.8, 0.9, 0.95 and 0.99. These simulations were based on the indicated alpha levels (1×10^-6^ [effectively Poisson], 0.2 and 0.5), 27 simulated cell types, 50 simulated cells per cell type, and 1000 simulated genes per theoretical Tau level.

### Tau p-value estimates

P-values for Tau being higher than predicted based on uniform expression across cell types were estimated by Monte Carlo simulations based on the actual number of cells profiled in each of the 27 cell types identified in the mid-gastrula *Ciona* dataset. Parameters for Poisson and negative binomial read count models were estimated by fitting Generalized Linear Models (GLMs) to the cell-level UMI counts using the Python statsmodels package. Total read counts per cell were used as model offsets to control for variable count depth. The statsmodels GLM tools cannot jointly fit mu and the overdispersion parameter alpha for negative binomial models, so we used scipy.optimize.minimize to find the alpha value that gives the model with highest log-likelihood for the two negative binomial models with individually fitted alpha values.

For the Poisson model, negative binomial common alpha model, and negative binomial fitted alpha model 2, we fit GLMs without using cell type as a categorical variable to give a single lambda or mu parameter per gene across all cell types. For negative binomial fitted alpha model 1 we first fit GLMs using cell type as a categorical variable to find the best alpha for each gene under the assumption that they are differentially expressed across cell types. We then fit new GLMs using those alpha values but no longer using cell type as a categorical variable to estimate the appropriate mu for uniform expression. The reasoning behind the two negative binomial fitted alpha models is that differential expression across cell types is likely to be a major cause of overdispersion. Model 2 estimates alpha without regard to distinct cell types and may thus provide an upper estimate on the likely range of alpha. Model 1 initially calculates alpha for each gene *with* regard to distinct cell types. Variable expression across cell types is thus accounted for in the separate mu values calculated for each cell type and alpha is expected to capture any remaining overdispersion resulting from latent cell states, bursty transcription, or the like. Model 1 rests on the assumption that this is a reasonable alpha level to simulate uniform expression across cell types. There is no strong reason to view model 1 as overly optimistic, but models 1 and 2 can be viewed as capturing a range of relevant alpha levels. In practice, all 4 models gave comparable results on this dataset but the differences would be increasingly important at higher read counts.

After fitting the model parameters, read counts were then simulated based on a single lambda or mu parameter for each gene across all cell types. Other model details (cell types, cells per cell type, total counts per cell) were directly based on the mid-gastrula scRNAseq dataset. Model mu and lambda parameters were calculated as counts per cell per total counts because of the offset for count depth. These were multiplied back out with count depth per cell to give rates per cell for the simulations. Each gene was simulated 25,00 times and p-values were estimated based on how often the simulated uniform expression Tau equaled or exceeded the observed Tau. These p-values were corrected for multiple comparisons using statsmodels.multipletests by the Benjamini and Hochberg False Discovery Rate method.

### Rate-cut tests

To estimate the sensitivity of observed Tau levels to expression level, we again simulated Tau using a negative binomial model and alpha values from fitted alpha model 1. Now, however, instead of using a uniform mu parameter across cell types we used the cell type-specific mu values estimated in the original GLM fits. These were cut in half to estimate the change in observed Tau for a 50% decrease in expression level. These half-rate simulated Taus were calculated for every gene but were binned in a two-dimensional grid for the streamline visualization in Fig. 5E-H.

### Gene Ontology enrichment analysis

Enriched GO codes for the ubiquitous, tissue-specific and highly tissue-specific gene sets discussed in the Results were identified using the GOATOOLS Python package [58], which is based on Fisher’s exact test and the hypergeometric distribution. The GO code annotation for *Ciona* KH2012 gene models was downloaded from the ANISEED ascidian community database [57]. It is based on homology to vertebrate orthologs and contains Molecular Function, Cellular Component and Biological Process codes. It required minor reformatting to be GOATOOLS compliant. The annotation is against ANISEED gene ids so we mapped those to the KH2012 transcript models used in this study with the appropriate ANISEED map file. We used the go-basic.obo ontology downloaded from geneontology.org. All comparisons used the set of all expressed genes in the mid-gastrula data set as the background set. P-values for enrichment or depletion were corrected for multiple comparisons using the Benjamini-Hochberg False Discovery Rate approach [51].

## Acknowledgements

We thank Wendy Reeves for comments on the manuscript.

## Author Contributions

CH and KP performed initial investigations of the Tau metric in the context of the mid-gastrula scRNAseq dataset. MV implemented most of the analyses reported here and wrote the first draft of the manuscript. MV, KP and CH revised the manuscript together.

